# Genomic epidemiology reveals antibiotic resistance transfer and polyclonal dissemination of *Acinetobacter baumannii* in a Paraguayan hospital

**DOI:** 10.1101/2025.04.03.647102

**Authors:** Elena Bello-Lopez, Anibal Kawabata, Jazmin Cantero, Sebastian Mendoza, Eduardo Pertile, Angeles Perez-Osegura, Miguel A. Cevallos, Humberto Peralta, Alejandro Aguilar-Vera, Santiago Castillo-Ramirez

## Abstract

*Acinetobacter baumannii* is a major nosocomial pathogen worldwide and, specifically, in Latin America. Genomic epidemiology has been instrumental in determining the transmission dynamics of *A. baumannii* in many countries of the world, yet some Latin American countries have conducted no genomic epidemiology studies. Here, we conduct the first genomic epidemiology study about this pathogen in Paraguay. We sequenced 43 isolates from a big tertiary hospital in Paraguay collected from different wards in 2021 and 2022. Our genomic epidemiology analyses, including almost 200 genomes and considering the main international clones (ICs), show that IC1, IC2, IC4, IC5 and IC7 were found in the hospital. We found novel genetic variation (3 novel Sequence Types as per the Oxford MLST scheme and one as per the Pasteur scheme) within IC7. Antibiotic susceptibility tests show that all but one of the Paraguayan isolates were resistant to carbapenems. Notably, 98% were classified as multidrug-resistant. We detected plasmids in almost all the Paraguayan isolates. Furthermore, we detected cases of recent horizontal transfer of important antibiotic-resistance genes between different ICs. On a general note, our findings highlight polyclonal spreading across different hospital wards and horizontal transfer of clinically relevant antibiotic-resistance genes among the different clones. On a more local note, this is the first genomic epidemiology study of *A. baumannii* in Paraguay and will be a reference point for future studies in the country and the region.

## Introduction

In many regions of the world antimicrobial resistance (AMR) is one of the major health issues. Latin America is no exception to this trend; on the contrary, many countries in this region are severely affected by AMR. A recent study estimated more than half a million deaths related to bacterial AMR in 35 countries from the Americas in 2019. Importantly, *Acinetobacter baumannii* was found among the top six bacterial pathogens (1). *A. baumannii* is a very relevant pathogen that very often is the cause of infections in critically ill patients in hospitals all over the world. In this respect, the World Health Organization considers carbapenem-resistant *A. baumannii* as one of the highest-priority bacteria for which novel therapies are required (2).

Genomic epidemiology has been extremely useful in understanding local and global patterns of dissemination of the main *A. baumannii* International Clones (ICs) in different parts of the world (3). Furthermore, in the last decade, genomic epidemiology studies have been conducted in several countries in Latin America to determine the dispersion of some ICs (4-9). For instance, recent studies have shown lineages belonging to IC2 and IC5 found in Brazilian and Mexican hospitals (7-10). Of note, IC2 is the most frequent IC worldwide, whereas IC5 is commonly found in Latin America. These genomic epidemiology studies have also helped study less well-known ICs. For instance, the much less studied IC7 was recently described in Bolivian hospitals (4) and also in a Brazilian hospital in Sao Paulo (11). These studies have also shown that multidrug-resistant (MDR) and even extremely resistant (XDR) isolates are commonplace in Latin America (6-8, 10). There is no doubt that genomic epidemiology has been instrumental in understanding the dissemination of *A. baumannii* and its resistance in Latin America. However, there are still some countries in the region for which no genomic epidemiology studies have yet been conducted, and thus, we do not know which ICs are disseminated in those countries.

Paraguay is one of those countries in Latin America for which genomic epidemiology of *A. baumannii* is lacking. In this regard, no information about the ICs circulating in this country has been generated yet. Considering this lack of knowledge, this study aimed to conduct the first genomic epidemiology in this country. We sequenced 43 isolates from a big tertiary hospital in Asuncion, Paraguay, collected from many different wards between 2021 and 2022. Our results show that several ICs are circulating in the hospital and most of these are MDR isolates. Furthermore, we noted cases of horizontal transfer of important antibiotic-resistance genes among the different clones.

## Methods

### Bacterial isolates, antibiotic susceptibility profiles

Isolates were collected from the tertiary trauma hospital “Dr. Manuel Giagni” in Asuncion, Paraguay, from 2021 to 2022. The list of the 43 isolates, along with their metadata (date of collection, isolation source, age and gender of the patient, etc.), is provided in Supplementary Table 1. Non-repetitive isolates were taken from routine patient samples with infections 48 hours after admission to the hospital. Initially, the bacteria were identified with VITEK^®^2 System bioMérieux. Antimicrobial susceptibility testing was performed by the agar disk diffusion method for 13 antibiotics (BD BBL™ Sensi-Disc™), described in the Clinical and Laboratory Standards Institute (CLSI) guidelines v.2024, these results were also added to Supplementary Table 1. Susceptibility to colistin was performed by broth microdilution at the Laboratorio Central de Salud Pública, Asunción, Paraguay, the breakpoints were resistant ≥4μg/mL and intermediate ≤2-0.25 μg/mL (CLSI, 2024). While tigecycline was evaluated by the Vitek2 System, it is known that this methodology has limitations, and there are no breakpoints for this antibiotic and only the MIC values obtained are indicated. The values for both antibiotics are reported in Supplementary Table 1.

### Genome sequencing and ST assignation

We sequenced 43 isolates. The DNA extraction was done using the ThermoScientific DNA Recovery kit. Twenty genomes were sequenced at BGI, Korea, using the DNB-Seq, which generates short reads. The other 23 genomes were sequenced at Novogene, Sacramento, US, employing the NovaSeq-6000 platform. Two genomes were also sequenced using Oxford Nanopore (MinIon). We validated the quality of the reads with FASTQC v0.12.1 and adapters were removed via Trimm-Galore v0.6.7. Genome assembly employed SPAdes v3.15.5 (12) and Unicycler v0.4.7 (13) for the hybrid assemblies. Genome assemblies were evaluated with Quast v5.2.0. Genomes were annotated using PROKKA v1.14.6 (14) and we confirmed the correct species assignation by running an Average Nucleotide Identity analysis where all the isolates showed higher than 95% ANI value. We conducted Sequence Type (ST) assignation as per both (Pasteur and Oxford) Multilocus Sequence Typing (MLST) schemes. This was done employing the PubMLST database (15).

### Phylogenies, *in silico* prediction of the resistome and horizontal gene transfer detection

To put the Paraguayan isolates in the context of the major ICs, we included a set of 148 genomes from IC1 to IC8. These genomes are listed in a previous work in supplementary table 1 (16). Thus, our first phylogenomic analysis included 191 genomes (the 43 Paraguayan isolates + 148 genomes from the ICs). With these genomes we constructed phylogeny based on single gene families without recombination, as we did in previous work (17). The phylogeny was constructed using RAxML (18). We used the CARD database version 3.2.7 (19) to conduct the *in silico* prediction of the resistome in the same that we did previously for other *A. baumannii* isolates (20) with the exception that coverage was set to 74% or higher. To detect very recent horizontal gene transfer events we proceeded as before (21). Briefly, we required that the allelic variants of a gene (or SNPs) were identical in the different ICs. This was done by conducting a global sequence alignment with the ggsearch36 program (22), requiring 100 identities and coverage between the allelic variants in the different clones.

### Plasmid detection and *in silico* prediction

The plasmid bands of the strains were visualized through Eckhardt’s technique (23). Cellular cultures were grown overnight in LB broth, diluted to 0.4 DO_620nm_ and centrifugated. Cells were gently resuspended with lysis solution (0.4 mg/ml RNAse, 1 mg/ml bromophenol blue in TE buffer, 40 *μ*L of lysozyme, 40 *μ*L of ficoll) and loaded onto a 0.75% agarose gel (pre-treated with 17 *μ*L SDS 10%/xylen cyanol in water per slot, under 100 V inverted current for 17 min). The final run was carried out at 40V for 2 h and then at 100V for 12 hours at normal current. The approximation of the molecular size of plasmid bands was calculated by comparing their band migration with those of plasmids of known size. A linear correlation was made in a semi-log graph. Previously reported strains were used for comparison: *A. haemolyticus* AN54 (plasmids of 45.4, 12.8, 11.4, 6.4 and 4.7 kb), *A. baumannii* ATCC17978 (149, 13.4 and 11.3 kb) and *E. coli* DH5 a/pTR102 (10.6 kb). Additionally, we also conducted an *in silico* prediction of plasmids on the Paraguayan isolates. Plasmids were inferred using the MOB-suite tool (24). Considering this *in silico* prediction, we took the circular plasmids and the incomplete plasmids but with a relaxase or replicase as more reliable inferences, as we did before for *A. junii* (25). For typing the potential plasmids, we considered both Bertini’s (26) and Hamidian’s (27) typing systems. The homologous genes (or proteins in the case of Bertni’s typing) were determined in the contigs of the analyzed strains via BLAST searches against the databases created by Hamidian and by Bertini with an identity ≥ 80% identity and ≥ 70% coverage.

## Results

### Epidemiology and genomic epidemiology of the Paraguayan isolates

This is a one-hospital, retrospective, observational study, and the isolates were collected in 2021 and 2022. We analyzed 43 isolates from hospitalized with signs of infections. The patients had stayed at least 48 hours in the hospital before the bacteria culture test was conducted. Supplementary Table 1 provides the epidemiological details of these isolates. Most of the isolates, 36 (84%), came from males, whereas just 7 (16%) came from women. The main source of isolates, with 21 isolates (49%), was tracheal secretions. The age range went from 6 to 91 years. The patients stayed in different wards (10 in total) but Emergency and Intensive Care Unit A (ICU-A) were the most frequent wards. Not unexpected, these two wards not only had most of the isolates but also were the most diverse in terms of the ICs found (read section below). The epidemiology data show that this is a diverse collection of isolates. To understand the genomic epidemiology of the Paraguayan isolates, we constructed a genome phylogeny (see methods). Importantly, we included 148 publicly available genomes to provide an adequate context regarding the major ICs. These genomes belong to the main ICs and we have used them in previous studies (16, 20). Our phylogeny shows two major trends (see Figure 1). First, we noted that several introductions of *A. baumannii* have occurred in the hospital. These can be easily noted as the Paraguayan isolates form several groups dispersed across the tree. Secondly, these Paraguayan groups are associated with different ICs. The most frequent IC was IC2, with 14 isolates within the clade. The distributions of the Paraguayan isolates within IC2, which were located within several IC2 sub-linages, imply different, independent introductions. The IC5 was the second most common, where 13 isolates were assigned to this IC. The third most frequent clone was IC7, having 12 isolates. As with IC2, it seems that several IC7 sub-linages have been introduced into the hospital. We noted an isolated (AbHTMGP-2689) that, although not within the IC7 clade, is very close to this IC. We also found three isolates belonging to IC1 and one isolate assigned to IC4. Although the Paraguayan isolates could be assigned to the main ICs, within those ICs some of the Paraguayan isolates represent novel STs not seen before. We discovered three novel STs as per the Oxford MLST scheme and one as per the Pasteur MLST scheme, which is ST2526 corresponding to isolates AbHTMGP-4890 and AbHTMGP-5204. Considering the Oxford MLST, the novel STs involved six isolates all related to IC7. These novel STs are: ST3274 (AbHTMGP-2689 isolate), ST3275 (2 isolates, AbHTMGP-4890 and AbHTMGP-5204) and ST3276 (3 isolates, AbHTMGP-863, AbHTMGP-949, and AbHTMGP-917). Collectively, these results show that *A. baumannii* has been introduced repeatedly in the hospital and that these introductions were from different ICs. Where the most frequent were IC2, IC5 and IC7. Yet, we also found novel genetic variation in IC7.

**Figure 1.**
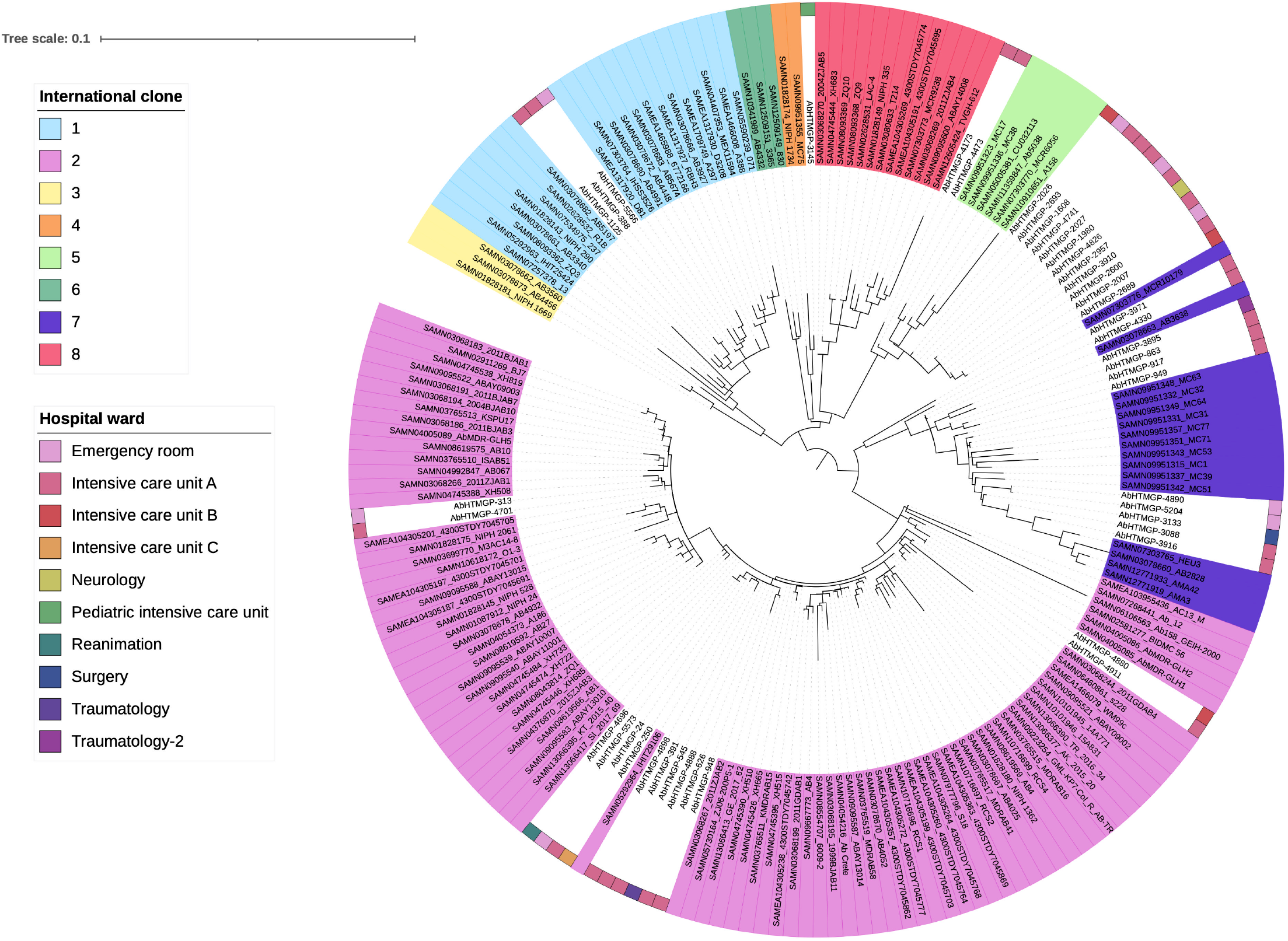
Phylogeny of the Paraguayan isolates plus the ICs. Phylogeny showing the relationships of the Paraguayan isolates with the main ICs. The different ICs are marked with different colours for the taxa labels; see color key. The wards where the isolates were collected are shown with coloured squares. The scale gives the number of substitutions per site.

### Two wards might be the source of dissemination of the ICs

Then, we look at the distribution of the ICs across the wards. Supplementary Figure 1 shows the different ICs found in the different wards. Two wards were not only the most frequent in terms of isolates but also had four different ICs. Intensive Care Unit A had 25 isolates, whereas Emergency had 8 isolates. Both wards had IC1, IC2, IC5 and IC7 in similar proportions. When one looks at the distributions of the wards, taking into account the ICs, in the phylogeny, It seems that the two previous wards could be the main source of dissemination for the other wards. For instance, the only isolate (AbHTMGP-2957) from Neurology is well within a clade (or group) of isolates from Emergency and IC-A, see IC5 and its surroundings in the phylogeny. Another example is isolate (AbHTMGP-3133) from Surgery, which also is in a group of isolates from Emergency and IC-A, see IC7 in the phylogeny. The exception seems to be the Pediatric Intensive Unit, which had the only isolate (AbHTMGP-3145) from IC4. Taken together, these results show that 2 wards are likely the main dissemination points in the hospital.

### High level of antibiotic resistance and presence of plasmids

Then we wanted to determine the level of resistance of these isolates. We noted that more than 98% of the isolates were MDR, as they were resistant to at least one antibiotic in at least 3 different antibiotic resistance classes (Figure 2). Notably, 93 to 98 % of the isolates were resistant to Ampicillin/Sulbactam, Piperacillin/Tazobactam, Cefepime, Cefotaxime, Ceftriaxone, Imipenem, Meropenem, Gentamicin, Ciprofloxacin, Levofloxacin. Furthermore, 74% were resistant to Ceftazidime, 86% to Amikacin, and 40% to Tetracycline. IC1 and IC2 were the only two ICs having isolates resistant to all these antibiotics (see Figure 2). However, all the five ICs have high levels of resistance. Of note, all isolates were categorized as colistin intermediate susceptible with MICs ranging from 0.25-2μg/mL. While for tigecycline, the following MICs were observed: 23% had 0.5-1 μg/mL, 53% had 2 μg/mL, 21% had 4 μg/mL, and 2% presented 8 μg/mL. In terms of the genetic determinants of antibiotic resistance, we found plenty.

**Figure 2.**
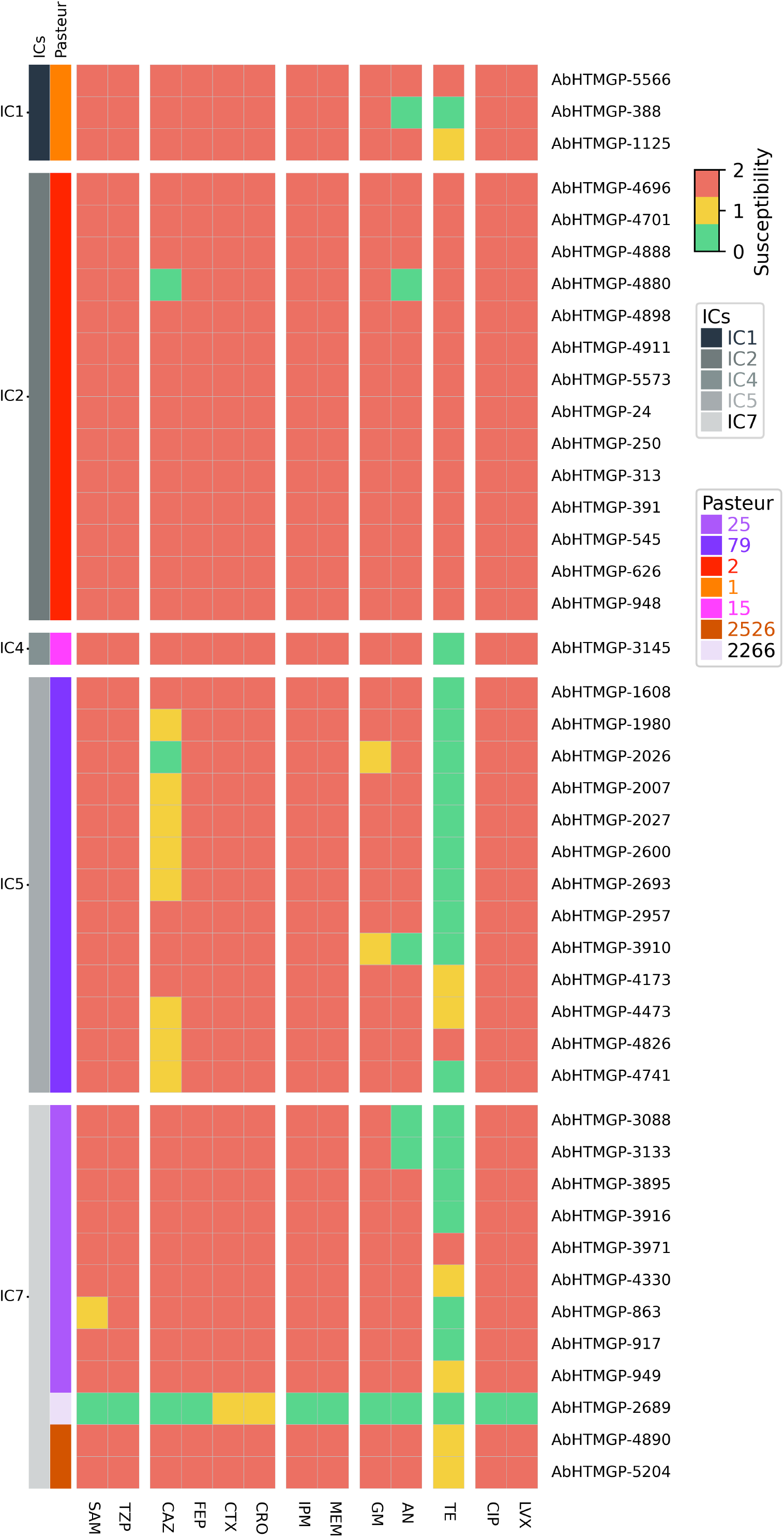
Antibiotic resistance profiles of the Paraguayan isolates. Antibiotic resistance profiles for the isolates. Green denotes susceptible, yellow intermediate, and salmon resistant. The vector colours on the left show the IC and ST (as per Pasteur MLST scheme) assignation.

Our *in silico* prediction of the resistome is shown in Figure 3. We noted that the isolates had several efflux pumps (RND, MSF, SMR and MATE) per genome; nonetheless, given that the over-expression of these pumps is what provides the resistance phenotypes, further expression analysis are required. We also found Tet (B) and Tet (C) pumps in isolates from IC2, IC5, and IC7. The beta-lactamase OXA-23 was present in all but one of the isolates and was the only acquired OXA detected. Of note, the only isolate (AbHTMGP-2689) not having OXA-23 was susceptible to all but one of the antibiotics tested (see Figure 2). Novel ADC beta-lactamase variants were identified and submitted to the NCBI. These are ADC-344 (accession number OR754316), present in IC1 isolates, and ADC-345 (OR753117), present in two IC7 isolates. Supplementary Figure 2 provides the genomic context of *armA*, and the aminoglycoside-modifying enzymes in the IC2 isolates. Considering resistance mediated by mutations, all but one isolate were resistant to Ciprofloxacin and Levofloxacin. Only AbHTMGP-2689 was susceptible, and this could be attributed to the fact that the S81L mutation was not found in *gyrA* and that it only had two point mutations (V104I, D105E) in *parC* compared to the other isolates. Interestingly, this isolate was the most susceptible to tall antibiotics tested and belongs to one (ST3274) of the novel STs as per the Oxford MLS scheme. Then, we determined the presence of plasmid bands in the Paraguayan isolates we ran Eckhardt plasmid profiles for 17 isolates. We made sure to have more isolates from the most abundant ICs. All 17 isolates had at least one plasmid (see Supplementary Figure 3). Whereas ten isolates (59%) had 2 or 3 plasmid bands, seven isolates (41%) had just one plasmid band. Of note, seven isolates had a big plasmid band (ca. 75-120 Kb), with one isolate (lane 14, Supplementary Figure 1) presenting two big plasmid bands (120 and 90 Kb, approx.). As expected, similar plasmid band profiles were found for isolates belonging to the same IC. Hence, plasmid band profiles show that plasmids are a common feature in these isolates. However, we fully acknowledge that Eckhardt’s technique does not allow discrimination between isoforms of the same plasmid, but it allows us to have an approximation of extrachromosomal elements carried by the isolates. Nonetheless, more analyses are required to sort this out. We also conducted an *in silico* prediction of plasmids. Considering only reliable inferences (see methods), we found 1.5 plasmids per isolate, on average (see Supplementary Table 2), which is not that different to what was found with the plasmid band profiles. We also typed the plasmids using two different strategies (see methods). We found that many plasmids could be assigned to different types (see Supplementary Table 2). Although both strategies typed a significant number of plasmids, more plasmids could be typed using Hamidian’s database. Considering the strategy employing Bertni’s database, we found 2 Rep groups (GR4 and G12) and just one Rep superfamily (Rep-3), see Supplementary Table 2. GR4 was the most frequent group. However, some of them cannot be assigned to any known type, and further studies are required to characterize them. Thus, both the plasmid band profiles and the *in silico* prediction show the presence of plasmids in the Paraguayan isolates. Finally, we noted that several ARGs were located in plasmids (see asterisks in Figure 3). Clear examples of this are the clinically relevant genes coding for OXA-23, TEM-1, Sul1, Sul2 and TetB. Collectively, these results show that Paraguayan isolates have important levels of resistance and frequent plasmid occurrence.

**Figure 3.**
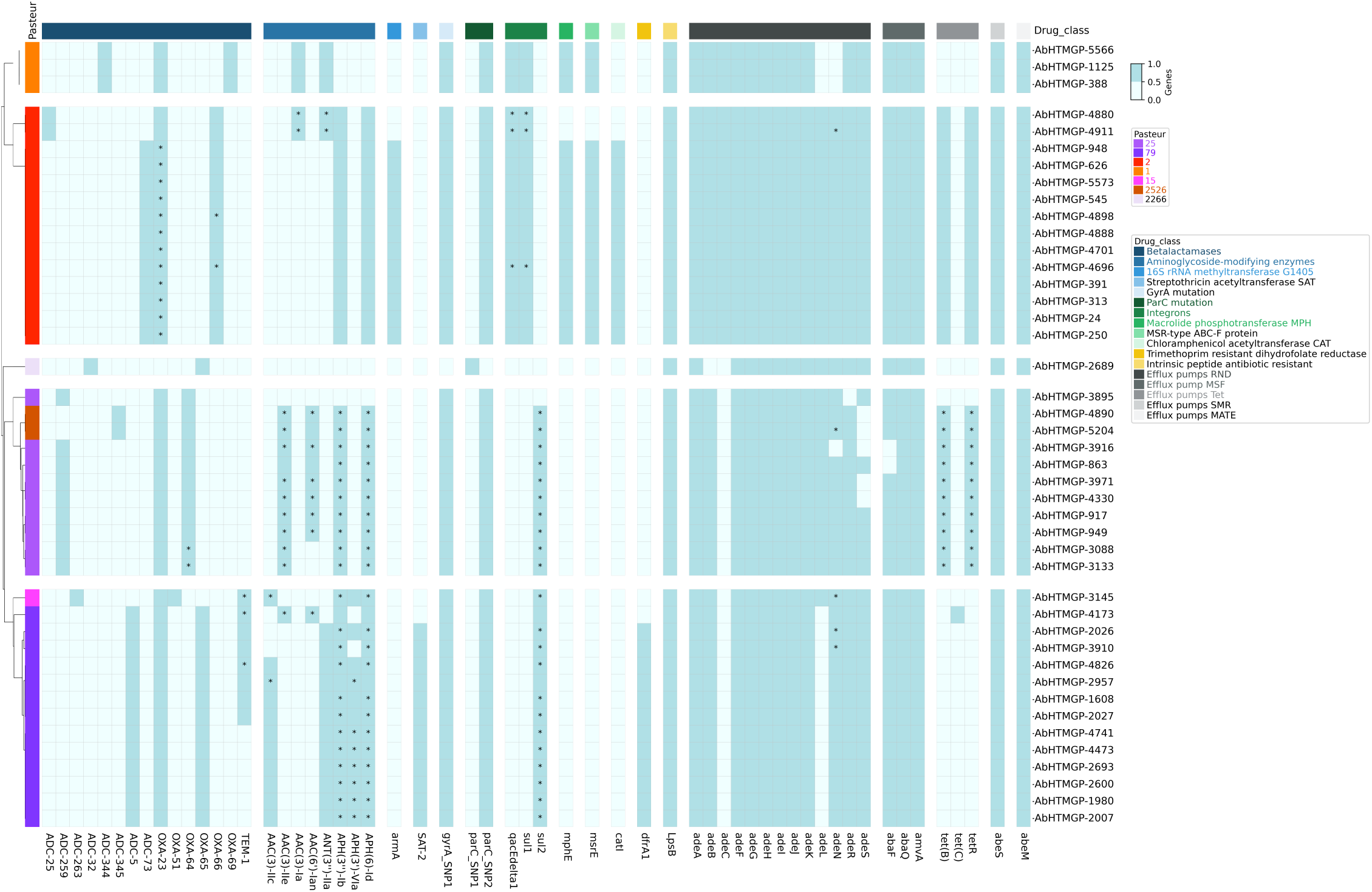
*In silico* prediction of the resistome of the Paraguayan isolates. Blue colour denotes the presence of the gene and light blue colour absence. The names of the ARGs are provided at the bottom. The vector colour on the right gives the ST as per the Pasteur MLST scheme. The asterisks show those ARGs that were predicted to be in plasmid signals.

### Horizontal antibiotic resistance gene transfer between the ICs

It is known that *A. baumannii* has an incredible ability to gain genes (28), in particular ARGs (21). Considering this ability and the co-existence of isolates from different ICs in the same wards, we wanted to know if there had been a transference of ARG among the ICs. We consider identical allelic variants as a hallmark of (very recent) horizontal gene transfer (HGT) events (see methods). Of note, if genes from different ICs exclusively experienced vertical transmission, they would exhibit several nucleotide changes in their sequences. Figure 4 shows the HGT events we found. We found that 19 ARGs that have endured HGT (see Figure 4). We noted clinically relevant ARGs, such as *bla*_OXA-23_ (OXA-23), *bla*_TEM-1_ (TEM-1) and AME genes. Some ICs share more HGT with other ICs. For instance, IC4 and IC7 are the ones showing the most HGT events with 7 ARGs. Followed by IC5 and IC7, which have 6 ARGs transferred. For their part, IC1 and IC2 present 7 ARGs. Thus, particular pairs of ICs are preferentially exchanging ARGs. Of note, IC1 seems to be the IC sharing fewer ARGs with the rest of the ICs; whereas, IC4 is involved in most HGT events. Whereas most genes have been transferred between just two ICs, only *bla*_OXA-23_ has been transferred among the four ICs. The second most transferred gene was the gene coding for APH(6)-ld, which was found in all the ICs but IC1.We noted the horizontal transfer of ARGs associated with acquired but also intrinsic resistance. Notably, in terms of the former, *bla*_OXA-23_ is the most salient case as this gene has been shuttled between all the ICs (see the dark purple lines in Figure 4). Considering the intrinsic resistance, we found that the genes coding for the adeIJK efflux pump have been transferred between some isolates of IC7 and IC4. Altogether, these results show that there have been some recent horizontal transfer of ARGs among the ICs within the hospital.

**Figure 4.**
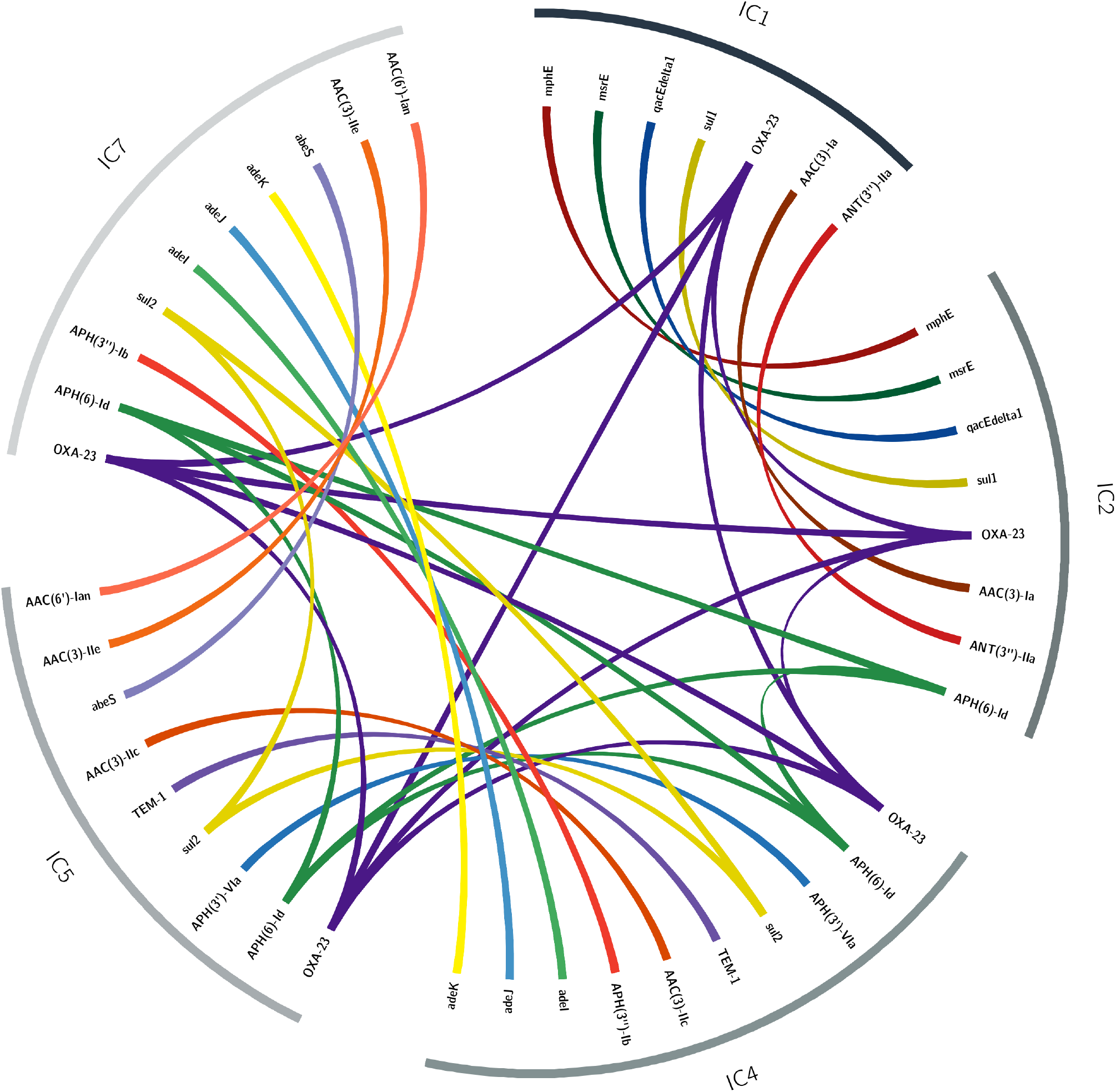
Horizontal gene transfer of ARGs. Circos plot showing the HGT events between ICs. The lines connecting the same gene in different ICs show the cases where identical allelic variants were found in different ICs. The colours of the lines denote different genes.

## Discussion

Genomic epidemiology of *A. baumannii* has been conducted in many parts of the world. This has been very instructive in understanding the dissemination of this pathogen in those regions. However, other areas of the world have been neglected. There are countries in Latin America for which very little if anything, is known about the molecular epidemiology of *A. baumannii*. Until this study, Paraguay was one of those countries. In this study, we did not only study the lineages circulating in a big tertiary hospital but also put them in a global context using genome representatives from the main ICs.

Our phylogenetic analysis shows that several introductions of *A. baumannii* into the hospital have occurred. These introductions were associated with five ICs. IC2 and IC5 were the most abundant clones in the hospital. This is not unexpected, as these ICs have been frequently described in other Latin American countries (7-10) and worldwide (3). However, many isolates were associated with the IC7, an IC not long ago reported in Bolivian hospitals (4). Of note, Bolivia and Paraguay are bordering countries. Considering this IC, we detected novel genetic variation: 3 novel Oxford STs in 6 isolates. It is worth mentioning that IC7 has not been studied as much as IC1, IC2, or IC3. We also found a single isolate belonging to IC4, an IC that has been described in Chile (29). To our knowledge, this is one of the first studies that found many ICs (5 clones) within one hospital in Latin America. Regarding the dissemination within the hospital, most of the isolates came from just two wards: Emergency and ICU-A. From those two wards, 77 % of the isolates were recovered. Furthermore, the phylogenetic tree suggests that these two wards were dissemination points to other wards. This highlights how certain wards (or places for that matter) could be major sources of infections within hospital settings. By the same token, this can be very helpful to infection control teams to pay special attention to these problematic places and vigorously decontaminate them.

We found high levels of resistance among the Paraguayan isolates. Almost all isolates were MDR and resistant to carbapenems (meropenem and imipenem). Although the isolates from all five ICs have a high amount of resistance, IC1 and IC2 were the only two clones where we found isolates resistant to almost all the antibiotics tested. Remarkably, we found that ICs are horizontally exchanging important ARGs. One of the most salient cases is *bla*_OXA-23_, which has been exchanged between all the 5 ICs. Notably, the HGT events were not restricted just to acquired ARGs but also included some classic examples of intrinsic resistance, such as the genes coding for RDN efflux pumps. We find that some pairs of ICs tend to preferentially exchange ARGs. This is not unexpected, and there might be several reasons for this. One could just be that those pairs tend to co-exist more often. Alternatively, those pairs might have more affinity in terms of the mobile genetic elements mobilizing the ARGs. Of note, our strategy to detect HGT events is biased toward very recent events. Thus, it underestimates the actual number of transfer events among the clones.

We acknowledge that our study has several limitations. The most relevant is that the isolates came from just one hospital and that the sampling was conducted for just 2 years. Nonetheless, given that there is no other study about the genomic epidemiology of *A. baumannii* in Paraguay, this study is highly relevant and it will be a point of reference for future studies in this country and its neighboring countries. In a broader context, our study highlights polyclonal dissemination across different wards in the same hospital and horizontal transfer of clinically relevant antibiotic-resistance genes among the different clones. Although, over the last few years, it has become clear that we need to pay attention to the non-human of *A. baumannii* (30-32), there are still some countries in the world for which not much information about their human clinical isolates is available. Thus, we also want to pay attention to those understudied countries to better understand the global epidemiology of *A. baumannii* (3).

## Data sharing statement

All the newly sequenced genomes have been deposited in the NCBI under BioProject PRJNA1012735. The publicly available genomes used are those described in a previous work in Supplementary Table 1 (16).

## Ethical approval

The protocol for this study was carefully reviewed and approved by the Hospital Ethical Committee [No. 04/2022 Acta 1/2022]. All isolates were collected during routine diagnosis and did not constitute additional risks to patients. Patient information was kept anonymous.

## Acknowledgements

We are grateful to Dra. Maria Eugenia León Ayala head of the section “Enfermedades Respiratorias y Meníngeas” at the “Laboratorio Central de Salud Pública”, Paraguay. We also thank Valeria Mateo-Estrada for submitting the new alleles to the PubMLST and helping with the visualization of Figure 1. We extend warm thanks to Victor Manuel del Moral Chaves and Alfredo J. Hernández-Alvarez for managing the servers and cluster where most of the *in silico* analyses were run. Finally, we extend many thanks to Laura Cervantes for helping with the plasmid profiles.

## SUPPLEMENTARY MATERIAL

**Supplementary Table 1**

The list of all the Paraguyan isolates sequenced. The table provides the metadata for each one of the isolates.

**Supplementary Table 2**

*In silico* prediction of plasmids. The table provides the prediction and typing of plasmids for each isolate.

**Supplementary Figure 1**

Isolates were sampled from different wards in the hospital. The colors in the bars show the IC designation for the isolates.

**Supplementary Figure 2**

Genomic context of armA, and AMEs in the IC2 isolates.

**Supplementary Figure 3**.

Plasmid band profiles of 17 Paraguayan isolates. Isolates are ordered as follows: Lane 1, AbHTMGP-5566. Lane 2, AbHTMGP-388. Lane 3, AbHTMGP-1125. Lane 4, AbHTMGP-4911. Lane 5, AbHTMGP-4696. Lane 6, AbHTMGP-4701. Lane 7, AbHTMGP-4888. Lane 8, AbHTMGP-3145. Lane 9, AbHTMGP-4173. Lane 10, AbHTMGP-2026. Lane 11, AbHTMGP-2957. Lane 12, AbHTMGP-2693. Lane 13, AbHTMGP-4330. Lane 14, AbHTMGP-863. Lane 15, AbHTMGP-4890. Lane 16, AbHTMGP-3916. Lane 17, AbHTMGP-2689. Lane 18, *A. haemolyticus* AN54. Lane 19, *A. baumannii* ATCC17978. Line 20, *E. coli* DH5a/pTR102.

