## Supplementary Material for "Genomic epidemiology reveals antibiotic resistance transfer and polyclonal dissemination of *Acinetobacter baumannii* in a Paraguayan hospital"

Supplementary Figure 1

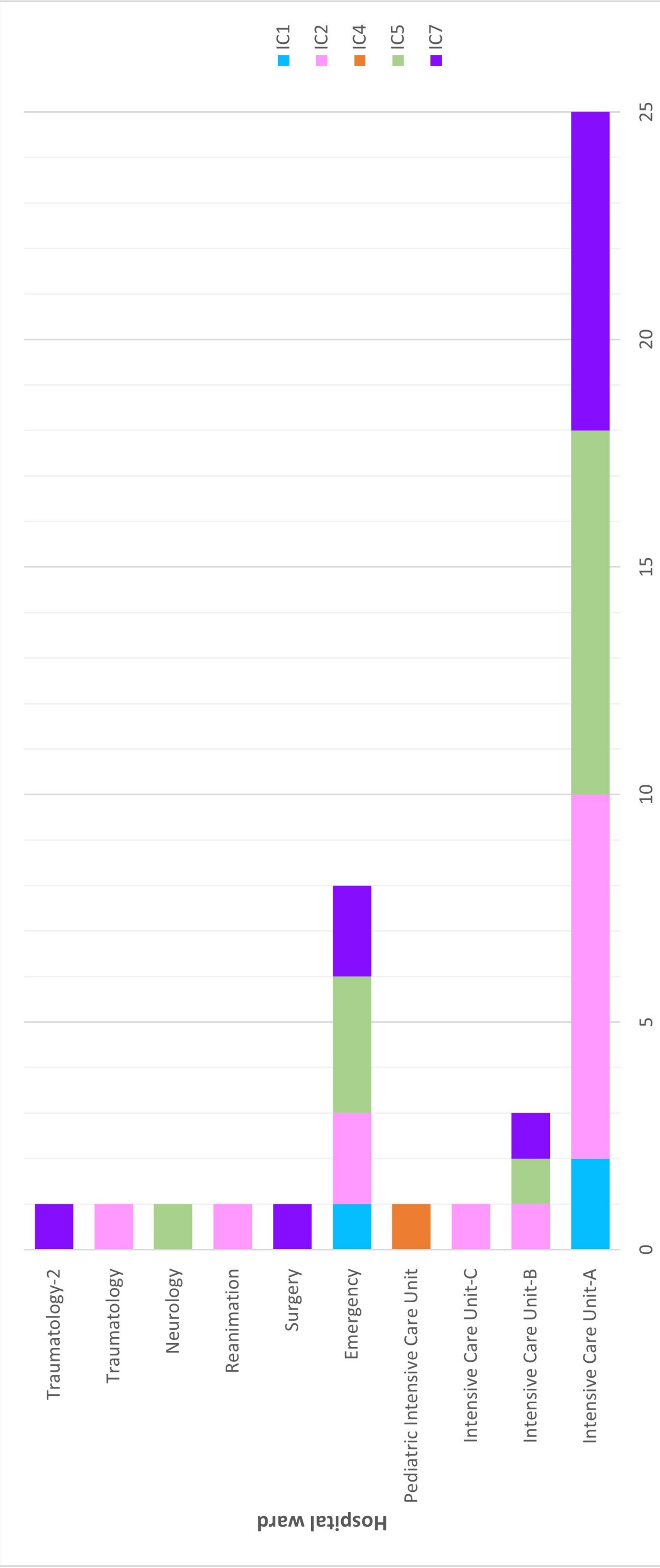


**Supplementary Figure 1**
**Isolates sampled from different wards in the hospital.**

Supplementary Figure 2

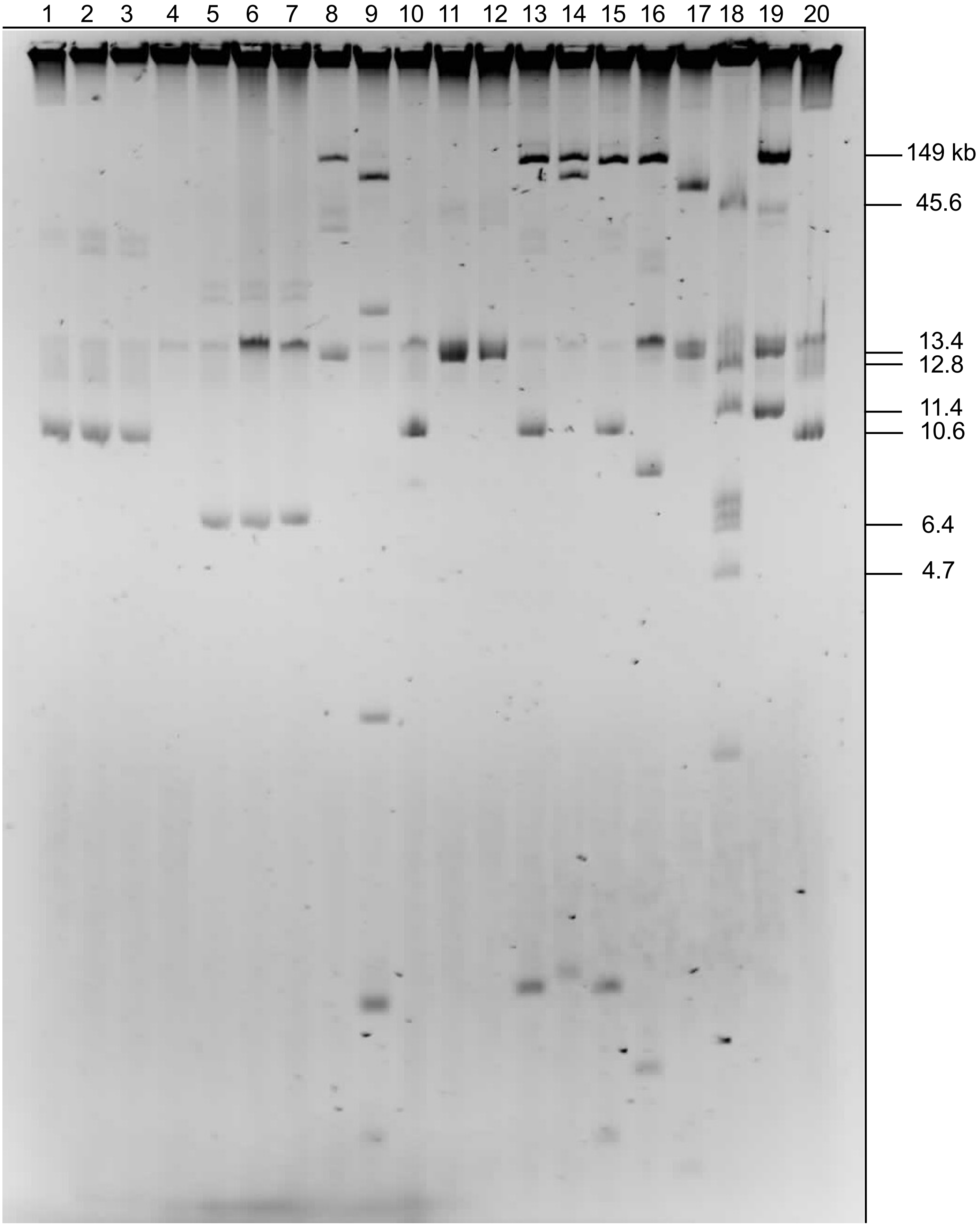

### Supplementary Table 1

**Supplementary Table 1**. The list of all the Paraguyan isolates sequenced. The table provides the metadata for each one of the isolates.

| NCBI | Date | Age | Gender | Ward | Source | ST (Oxford/Pasteur) | International Clone | Betalactamases |
|---|---|---|---|---|---|---|---|---|
| AbHTMGP-1608 | 17/03/21 | 21 | m | Intensive care unit A | Tracheal secretion | 1283/79 | IC5 | OXA-65, OXA-23, ADC-5, TEM-1 |
| AbHTMGP-1980 | 05/04/21 | 39 | m | Emergency room | Tracheal secretion | 1283/79 | IC5 | OXA-65, OXA-23, ADC-5 |
| AbHTMGP-2026 | 07/04/21 | 28 | m | Intensive care unit B | Tracheal secretion | 1283/79 | IC5 | OXA-65, OXA-23, ADC-5, TEM-1 |
| AbHTMGP-2007 | 07/04/21 | 48 | m | Intensive care unit A | Blood culture | 1283/79 | IC5 | OXA-65, OXA-23,ADC-5 |
| AbHTMGP-2027 | 08/04/21 | 28 | m | Intensive care unit A | Blood culture | 1283/79 | IC5 | OXA-65, OXA-23, ADC-5, TEM-1 |
| AbHTMGP-2600 | 07/05/21 | 33 | m | Intensive care unit A | Blood culture | 1283/79 | IC5 | OXA-65,OXA-23, ADC-5 |
| AbHTMGP-2689 | 12/05/21 | 21 | m | Intensive care unit B | Tracheal secretion | 3274/2266 | IC7 | OXA-65, ADC-32 |
| AbHTMGP-2693 | 12/05/21 | 60 | m | Emergency room | Tracheal secretion | 1283/79 | IC5 | OXA-65, OXA-23, ADC-5 |
| AbHTMGP-2957 | 26/05/21 | 32 | f | Neurology | Purulent discharge | 1283/79 | IC5 | OXA-65, OXA-23, ADC-5, TEM-1 |
| AbHTMGP-3088 | 01/06/21 | 26 | m | Intensive care unit A | Tracheal secretion | 1489/25 | IC7 | OXA-64, OXA-23, ADC-259 |
| AbHTMGP-3133 | 04/06/21 | 56 | m | Surgery | Urine | 1489/25 | IC7 | OXA-64, OXA-23, ADC-259 |
| AbHTMGP-3145 | 04/06/21 | 6 | m | Pediatric intensive care unit | Tracheal secretion | 236/15 | IC4 | OXA-51, OXA-23, ADC-263, TEM-1 |
| AbHTMGP-3895 | 13/07/21 | 47 | f | Traumatology-2 | Purulent discharge | 691/25 | IC7 | OXA-64, OXA-23, ADC-259 |
| AbHTMGP-3910 | 13/07/21 | 19 | m | Intensive care unit A | Tracheal secretion | 1283/79 | IC5 | OXA-65, OXA-23, ADC-5, TEM-1 |
| AbHTMGP-3916 | 13/07/21 | 21 | m | Intensive care unit A | Blood culture | 490/25 | IC7 | OXA-64, OXA-23, ADC-259 |
| AbHTMGP-3971 | 15/07/21 | 22 | m | Intensive care unit A | Tracheal secretion | 691/25 | IC7 | OXA-64, OXA-23, ADC-259 |
| AbHTMGP-4173 | 27/07/21 | 28 | m | Intensive care unit A | Blood culture | 758/79 | IC5 | OXA-65, OXA-23, ADC-5, TEM-1 |
| AbHTMGP-4330 | 04/08/21 | 49 | f | Intensive care unit A | Purulent discharge | 1489/25 | IC7 | OXA-64, OXA-23, ADC-259 |
| AbHTMGP-4473 | 11/08/21 | 21 | m | Intensive care unit A | Tracheal secretion | 1283/79 | IC5 | OXA-65, OXA-23, ADC-5 |
| AbHTMGP-4696 | 18/08/21 | 27 | m | Reanimation | Tracheal secretion | 195/2 | IC2 | OXA-66, OXA-23, ADC-73 |
| AbHTMGP-4701 | 18/08/21 | 31 | m | Intensive care unit A | Tracheal secretion | 195/2 | IC2 | OXA-66, OXA-23, ADC-73 |
| AbHTMGP-4826 | 19/08/21 | 21 | m | Intensive care unit A | Tracheal secretion | 1283/79 | IC5 | OXA-65, OXA-23, ADC-5, TEM-1 |
| AbHTMGP-4741 | 20/08/21 | 18 | m | Intensive care unit A | Tracheal secretion | 1283/79 | IC5 | OXA-65, OXA-23, ADC-5 |
| AbHTMGP-4890 | 26/08/21 | 91 | m | Emergency room | Purulent discharge | 3275/2526 | IC7 | OXA-64, OXA-23, ADC-345 |
| AbHTMGP-4888 | 26/08/21 | 30 | f | Traumatology | Purulent discharge | 195/2 | IC2 | OXA-66, OXA-23, ADC-73 |
| AbHTMGP-4880 | 26/08/21 | 34 | m | Intensive care unit B | Tracheal secretion | 208/2 | IC2 | OXA-66, OXA-23, ADC-25 |
| AbHTMGP-4898 | 27/08/21 | 77 | m | Intensive care unit A | Urine | 195/2 | IC2 | OXA-66, OXA-23, ADC-73 |
| AbHTMGP-4911 | 27/08/21 | 22 | m | Intensive care unit A | Bronchoalveolar lavage | 208/2 | IC2 | OXA-66, OXA-23, ADC-25 |
| AbHTMGP-5204 | 10/09/21 | 19 | m | Emergency room | Blood culture | 3275/2526 | IC7 | OXA-64, OXA-23, ADC-345 |
| AbHTMGP-5573 | 24/09/21 | 33 | m | Emergency room | Blood culture | 195/2 | IC2 | OXA-66, OXA-23, ADC-73 |
| AbHTMGP-5566 | 24/09/21 | 51 | f | Emergency room | Purulent discharge | 231/1 | IC1 | OXA-69, OXA-23, ADC-344 |
| AbHTMGP-24 | 02/01/22 | 48 | m | Intensive care unit A | Tracheal secretion | 195/2 | IC2 | OXA-66, OXA-23,ADC-73 |
| AbHTMGP-250 | 12/01/22 | 39 | m | Intensive care unit C | Tracheal secretion | 195/2 | IC2 | OXA-66, OXA-23, ADC-73 |
| AbHTMGP-313 | 16/01/22 | 41 | m | Emergency room | Purulent discharge | 195/2 | IC2 | OXA-66, OXA-23, ADC-73 |
| AbHTMGP-391 | 19/01/22 | 29 | m | Intensive care unit A | Urine | 195/2 | IC2 | OXA-66, OXA-23, ADC-73 |
| AbHTMGP-388 | 19/01/22 | 28 | m | Intensive care unit A | Catheter tip | 231/1 | IC1 | OXA-69, OXA-23, ADC-344 |
| AbHTMGP-545 | 26/01/22 | 23 | m | Intensive care unit A | Tracheal secretion | 195/2 | IC2 | OXA-66, OXA-23,ADC-73 |
| AbHTMGP-626 | 30/01/22 | 31 | m | Intensive care unit A | Tracheal secretion | 195/2 | IC2 | OXA-66, OXA-23, ADC-73 |
| AbHTMGP-863 | 09/02/22 | 22 | f | Intensive care unit A | Blood culture | 3276/25 | IC7 | OXA-64, OXA-23, ADC-259 |
| AbHTMGP-917 | 11/02/22 | 21 | f | Intensive care unit A | Tracheal secretion | 3276/25 | IC7 | OXA-64, OXA-23,ADC-259 |
| AbHTMGP-948 | 13/02/22 | 37 | m | Intensive care unit A | Tracheal secretion | 195/2 | IC2 | OXA-66, OXA-23, ADC-73 |
| AbHTMGP-949 | 14/02/22 | 27 | m | Intensive care unit A | Urine | 3276/25 | IC7 | OXA-64, OXA-23, ADC-259 |
| AbHTMGP-1125 | 21/02/22 | 25 | m | Intensive care unit A | Blood culture | 231/1 | IC1 | OXA-69, OXA-23, ADC-344 |

| NCBI | Aminoglucoside modifying enzymes | methyltransferas | otothricin acetyltransferase ( | GyrA mutation | ParC mutation | Integrons elements | RND Efflux pumps |
|---|---|---|---|---|---|---|---|
| AbHTMGP-1608 | AAC(3)-IIc, APH(3'')-Ib,APH(3')-VIa, APH(6)-Id, ANT(3'')-IIa | | SAT-2 | S81L | S84L, V104I, D105E | sul2 | adeA, adeB, adeF, adeG, adeH, adeI, adeJ, adeK, adeN, adeR, adeS |
| AbHTMGP-1980 | AAC(3)-IIc, APH(3'')-Ib,APH(3')-VIa, APH(6)-Id, ANT(3'')-IIa | | SAT-2 | S81L | S84L, V104I, D105E | sul2 | adeA, adeB, adeF, adeG, adeH, adeI, adeJ, adeK, adeN, adeR, adeS |
| AbHTMGP-2026 | APH(3'')-Ib, APH(6)-Id, APH(3')-VIa, ANT(3'')-IIa | | SAT-2 | S81L | S84L, V104I, D105E | sul2 | adeA, adeB, adeF, adeG, adeH, adeI, adeJ, adeK, adeN, adeR, adeS |
| AbHTMGP-2007 | AAC(3)-IIc, APH(3'')-Ib,APH(3')-VIa, APH(6)-Id, ANT(3'')-IIa | | SAT-2 | S81L | S84L, V104I, D105E | sul2 | adeA, adeB, adeF, adeG, adeH, adeI, adeJ, adeK, adeN, adeR, adeS |
| AbHTMGP-2027 | AAC(3)-IIc, APH(3'')-Ib,APH(3')-VIa, APH(6)-Id, ANT(3'')-IIa | | SAT-2 | S81L | S84L, V104I, D105E | sul2 | adeA, adeB, adeF, adeG, adeH, adeI, adeJ, adeK, adeN, adeR, adeS |
| AbHTMGP-2600 | AAC(3)-IIc, APH(3'')-Ib,APH(3')-VIa, APH(6)-Id, ANT(3'')-IIa | | SAT-2 | S81L | S84L, V104I, D105E | sul2 | adeA, adeB, adeF, adeG, adeH, adeI, adeJ, adeK, adeN, adeR, adeS |
| AbHTMGP-2689 | | | | | V104I, D105E | | adeA, adeF, adeG, adeH, adeI, adeJ, adeK, adeL, adeN, adeR, adeS |
| AbHTMGP-2693 | AAC(3)-IIc, APH(3'')-Ib,APH(3')-VIa, APH(6)-Id, ANT(3'')-IIa | | SAT-2 | S81L | S84L, V104I, D105E | sul2 | adeA, adeB, adeF, adeG, adeH, adeI, adeJ, adeK, adeN, adeR, adeS |
| AbHTMGP-2957 | AAC(3)-IIc, APH(3'')-Ib,APH(3')-VIa, APH(6)-Id, ANT(3'')-IIa | | SAT-2 | S81L | S84L, V104I, D105E | sul2 | adeA, adeB, adeF, adeG, adeH, adeI, adeJ, adeK, adeN, adeR, adeS |
| AbHTMGP-3088 | AAC(3)-IIe, APH(3'')-Ib, APH(6)-Id | | | S81L | S84L, V104I, D105E | sul2 | adeA, adeB, adeF, adeG, adeH, adeI, adeJ, adeK, adeL, adeN, adeR, adeS |
| AbHTMGP-3133 | AAC(3)-IIe, APH(3'')-Ib, APH(6)-Id | | | S81L | S84L, V104I, D105E | sul2 | adeA, adeB, adeF, adeG, adeH, adeI, adeJ, adeK, adeL, adeN, adeR, adeS |
| AbHTMGP-3145 | AAC(3)-IIc, APH(3'')-Ib,APH(3')-VIa, APH(6)-Id | | | S81L | S84L, V104I, D105E | sul2 | adeA, adeB, adeF, adeG, adeH, adeI, adeJ, adeK, adeL, adeN, adeR, adeS |
| AbHTMGP-3895 | | | | S81L | S84L, V104I, D105E | | adeA, adeB, adeF, adeG, adeH, adeI, adeJ, adeK, adeL, adeN, adeS |
| AbHTMGP-3910 | APH(3'')-Ib, APH(6)-Id, ANT(3'')-IIa | | SAT-2 | S81L | S84L, V104I, D105E | sul2 | adeA, adeB, adeF, adeG, adeH, adeI, adeJ, adeK, adeN, adeR, adeS |
| AbHTMGP-3916 | AAC(3)-IIe, AAC(6')-Ian, APH(3'')-Ib, APH(6)-Id | | | S81L | S84L, V104I, D105E | sul2 | adeA, adeB, adeF, adeG, adeH, adeI, adeJ, adeK, adeL, adeR |
| AbHTMGP-3971 | AAC(3)-IIe, AAC(6')-Ian, APH(3'')-Ib, APH(6)-Id | | | S81L | S84L, V104I, D105E | sul2 | adeA, adeB, adeF, adeG, adeH, adeI, adeJ, adeK, adeL, adeN, adeR |
| AbHTMGP-4173 | APH(3'')-Ib, APH(6)-Id, AAC(3)-IIe, AAC(6')-Ian | | | S81L | S84L, V104I, D105E | sul2 | adeA, adeB, adeF, adeG, adeH, adeI, adeJ, adeK, adeN, adeR, adeS |
| AbHTMGP-4330 | AAC(3)-IIe, AAC(6')-Ian, APH(3'')-Ib, APH(6)-Id | | | S81L | S84L, V104I, D105E | sul2 | adeA, adeB, adeF, adeG, adeH, adeI, adeJ, adeK, adeL, adeN, adeR |
| AbHTMGP-4473 | AAC(3)-IIc, APH(3'')-Ib,APH(3')-VIa, APH(6)-Id, ANT(3'')-IIa | | SAT-2 | S81L | S84L, V104I, D105E | sul2 | adeA, adeB, adeF, adeG, adeH, adeI, adeJ, adeK, adeN, adeR, adeS |
| AbHTMGP-4696 | APH(3'')-Ib, APH(6)-Id | armA | | S81L | S84L, V104I, D105E | qacEdelta1, sul1 | adeA, adeB, adeC, adeF, adeG, adeH, adeI, adeJ, adeK, adeL, adeN, adeR, adeS |
| AbHTMGP-4701 | APH(3'')-Ib, APH(6)-Id | armA | | S81L | S84L, V104I, D105E | qacEdelta1, sul1 | adeA, adeB, adeC, adeF, adeG, adeH, adeI, adeJ, adeK, adeL, adeN, adeR, adeS |
| AbHTMGP-4826 | AAC(3)-IIc, APH(3'')-Ib,APH(3')-VIa, APH(6)-Id, ANT(3'')-IIa | | SAT-2 | S81L | S84L, V104I, D105E | sul2 | adeA, adeB, adeF, adeG, adeH, adeI, adeJ, adeK, adeN, adeR, adeS |
| AbHTMGP-4741 | AAC(3)-IIc, APH(3'')-Ib,APH(3')-VIa, APH(6)-Id, ANT(3'')-IIa | | SAT-2 | S81L | S84L, V104I, D105E | sul2 | adeA, adeB, adeF, adeG, adeH, adeI, adeJ, adeK, adeN, adeR, adeS |
| AbHTMGP-4890 | AAC(3)-IIe, AAC(6')-Ian, APH(3'')-Ib, APH(6)-Id | | | S81L | S84L, V104I, D105E | sul2 | adeA, adeB, adeF, adeG, adeH, adeI, adeJ, adeK, adeL, adeN, adeR |
| AbHTMGP-4888 | APH(3'')-Ib, APH(6)-Id | armA | | S81L | S84L, V104I, D105E | qacEdelta1, sul1 | adeA, adeB, adeC, adeF, adeG, adeH, adeI, adeJ, adeK, adeL, adeN, adeR, adeS |
| AbHTMGP-4880 | AAC(3)-Ia, APH(3'')-Ib, APH(6)-Id, ANT(3'')-IIa | | | S81L | S84L, V104I, D105E | qacEdelta1, sul1 | adeA, adeB, adeC, adeF, adeG, adeH, adeI, adeJ, adeK, adeL, adeN, adeR, adeS |
| AbHTMGP-4898 | APH(3'')-Ib, APH(6)-Id | armA | | S81L | S84L, V104I, D105E | qacEdelta1, sul1 | adeA, adeB, adeC, adeF, adeG, adeH, adeI, adeJ, adeK, adeL, adeN, adeR, adeS |
| AbHTMGP-4911 | AAC(3)-Ia, APH(3'')-Ib, APH(6)-Id, ANT(3'')-IIa | | | S81L | S84L, V104I, D105E | qacEdelta1, sul1 | adeA, adeB, adeC, adeF, adeG, adeH, adeI, adeJ, adeK, adeL, adeN, adeR, adeS |
| AbHTMGP-5204 | AAC(3)-IIe, AAC(6')-Ian, APH(3'')-Ib, APH(6)-Id | | | S81L | S84L, V104I, D105E | sul2 | adeA, adeB, adeF, adeG, adeH, adeI, adeJ, adeK, adeL, adeN, adeR |
| AbHTMGP-5573 | APH(3'')-Ib, APH(6)-Id | armA | | S81L | S84L, V104I, D105E | qacEdelta1, sul1 | adeA, adeB, adeC, adeF, adeG, adeH, adeI, adeJ, adeK, adeL, adeN, adeR, adeS |
| AbHTMGP-5566 | AAC(3)-Ia, ANT(3'')-IIa | | | S81L | S84L, V104I, D105E | qacEdelta1, sul1 | adeA, adeB, adeC, adeF, adeG, adeH, adeI, adeJ, adeK, adeR, adeS |
| AbHTMGP-24 | APH(3'')-Ib, APH(6)-Id | armA | | S81L | S84L, V104I, D105E | qacEdelta1, sul1 | adeA, adeB, adeC, adeF, adeG, adeH, adeI, adeJ, adeK, adeL, adeN, adeR, adeS |
| AbHTMGP-250 | APH(3'')-Ib, APH(6)-Id | armA | | S81L | S84L, V104I, D105E | qacEdelta1, sul1 | adeA, adeB, adeC, adeF, adeG, adeH, adeI, adeJ, adeK, adeL, adeN, adeR, adeS |
| AbHTMGP-313 | APH(3'')-Ib, APH(6)-Id | armA | | S81L | S84L, V104I, D105E | qacEdelta1, sul1 | adeA, adeB, adeC, adeF, adeG, adeH, adeI, adeJ, adeK, adeL, adeN, adeR, adeS |
| AbHTMGP-391 | APH(3'')-Ib, APH(6)-Id | armA | | S81L | S84L, V104I, D105E | qacEdelta1, sul1 | adeA, adeB, adeC, adeF, adeG, adeH, adeI, adeJ, adeK, adeL, adeN, adeR, adeS |
| AbHTMGP-388 | AAC(3)-Ia, ANT(3'')-IIa | | | S81L | S84L, V104I, D105E | qacEdelta1, sul1 | adeA, adeB, adeC, adeF, adeG, adeH, adeI, adeJ, adeK, adeR, adeS |
| AbHTMGP-545 | APH(3'')-Ib, APH(6)-Id | armA | | S81L | S84L, V104I, D105E | qacEdelta1, sul1 | adeA, adeB, adeC, adeF, adeG, adeH, adeI, adeJ, adeK, adeL, adeN, adeR, adeS |
| AbHTMGP-626 | APH(3'')-Ib, APH(6)-Id | armA | | S81L | S84L, V104I, D105E | qacEdelta1, sul1 | adeA, adeB, adeC, adeF, adeG, adeH, adeI, adeJ, adeK, adeL, adeN, adeR, adeS |
| AbHTMGP-863 | AAC(3)-IIe, AAC(6')-Ian, APH(3'')-Ib, APH(6)-Id | | | S81L | S84L, V104I, D105E | sul2 | adeA, adeB, adeF, adeG, adeH, adeI, adeJ, adeK, adeL, adeN, adeR, adeS |
| AbHTMGP-917 | AAC(3)-IIe, AAC(6')-Ian, APH(3'')-Ib, APH(6)-Id | | | S81L | S84L, V104I, D105E | sul2 | adeA, adeB, adeF, adeG, adeH, adeI, adeJ, adeK, adeL, adeN, adeR, adeS |
| AbHTMGP-948 | APH(3'')-Ib, APH(6)-Id | armA | | S81L | S84L, V104I, D105E | qacEdelta1, sul1 | adeA, adeB, adeC, adeF, adeG, adeH, adeI, adeJ, adeK, adeL, adeN, adeR, adeS |
| AbHTMGP-949 | AAC(3)-IIe, APH(3'')-Ib, APH(6)-Id,AAC(6')-Ian | | | S81L | S84L, V104I, D105E | sul2 | adeA, adeB, adeF, adeG, adeH, adeI, adeJ, adeK, adeL, adeN, adeR, adeS |
| AbHTMGP-1125 | AAC(3)-Ia, ANT(3'')-IIa | | | S81L | S84L, V104I, D105E | qacEdelta1, sul1 | adeA, adeB, adeC, adeF, adeG, adeH, adeI, adeJ, adeK, adeR, adeS |

| NCBI | MSF Efflux pumps | et Efflux pump | SMR Efflux pumps | MATE Efflux pumps | Macrolide phosphotransferase (MPH) | msr-type ABC-F protein | nicol acetyltransferase | im resistant dihydrofolate re | nic peptide antibiotic resistan | R or PDR Cla |
|---|---|---|---|---|---|---|---|---|---|---|
| AbHTMGP-1608 | AbaF, AbaQ, AmvA | | abeS | abeM | | | | dfrA1 | LpsB | MDR |
| AbHTMGP-1980 | AbaF, AbaQ, AmvA | | abeS | abeM | | | | dfrA1 | LpsB | MDR |
| AbHTMGP-2026 | AbaF, AbaQ, AmvA | | abeS | abeM | | | | dfrA1 | LpsB | MDR |
| AbHTMGP-2007 | AbaF, AbaQ, AmvA | | abeS | abeM | | | | dfrA1 | LpsB | MDR |
| AbHTMGP-2027 | AbaF, AbaQ, AmvA | | abeS | abeM | | | | dfrA1 | LpsB | MDR |
| AbHTMGP-2600 | AbaF, AbaQ, AmvA | | abeS | abeM | | | | dfrA1 | LpsB | MDR |
| AbHTMGP-2689 | AbaF, AbaQ, AmvA | | abeS | abeM | | | | | LpsB | NC |
| AbHTMGP-2693 | AbaF, AbaQ, AmvA | | abeS | abeM | | | | dfrA1 | LpsB | MDR |
| AbHTMGP-2957 | AbaF, AbaQ, AmvA | | abeS | abeM | | | | dfrA1 | LpsB | MDR |
| AbHTMGP-3088 | AbaF, AbaQ, AmvA | tet(B), tetR | abeS | abeM | | | | | LpsB | MDR |
| AbHTMGP-3133 | AbaF, AbaQ, AmvA | tet(B), tetR | abeS | abeM | | | | | LpsB | MDR |
| AbHTMGP-3145 | AbaF, AbaQ, AmvA | | abeS | abeM | | | | | LpsB | MDR |
| AbHTMGP-3895 | AbaF, AbaQ, AmvA | | abeS | abeM | | | | | LpsB | MDR |
| AbHTMGP-3910 | AbaF, AbaQ, AmvA | | abeS | abeM | | | | dfrA1 | LpsB | MDR |
| AbHTMGP-3916 | AbaQ, AmvA | tet(B), tetR | abeS | abeM | | | | | LpsB | MDR |
| AbHTMGP-3971 | AbaF, AbaQ, AmvA | tet(B), tetR | abeS | abeM | | | | | LpsB | MDR |
| AbHTMGP-4173 | AbaF, AbaQ, AmvA | tet(C) | abeS | abeM | | | | | LpsB | MDR |
| AbHTMGP-4330 | AbaF, AbaQ, AmvA | tet(B), tetR | abeS | abeM | | | | | LpsB | MDR |
| AbHTMGP-4473 | AbaF, AbaQ, AmvA | | abeS | abeM | | | | dfrA1 | LpsB | MDR |
| AbHTMGP-4696 | AbaF, AbaQ, AmvA | tet(B), tetR | abeS | abeM | mphE | msrE | catI | | LpsB | MDR |
| AbHTMGP-4701 | AbaF, AbaQ, AmvA | tet(B), tetR | abeS | abeM | mphE | msrE | catI | | LpsB | MDR |
| AbHTMGP-4826 | AbaF, AbaQ, AmvA | | abeS | abeM | | | | dfrA1 | LpsB | MDR |
| AbHTMGP-4741 | AbaF, AbaQ, AmvA | | abeS | abeM | | | | dfrA1 | LpsB | MDR |
| AbHTMGP-4890 | AbaF, AbaQ, AmvA | tet(B), tetR | abeS | abeM | | | | | LpsB | MDR |
| AbHTMGP-4888 | AbaF, AbaQ, AmvA | tet(B), tetR | abeS | abeM | mphE | msrE | catI | | LpsB | MDR |
| AbHTMGP-4880 | AbaF, AbaQ, AmvA | tet(B), tetR | abeS | abeM | | | | | LpsB | MDR |
| AbHTMGP-4898 | AbaF, AbaQ, AmvA | tet(B), tetR | abeS | abeM | mphE | msrE | catI | | LpsB | MDR |
| AbHTMGP-4911 | AbaF, AbaQ, AmvA | tet(B), tetR | abeS | abeM | | | | | LpsB | MDR |
| AbHTMGP-5204 | AbaF, AbaQ, AmvA | tet(B), tetR | abeS | abeM | | | | | LpsB | MDR |
| AbHTMGP-5573 | AbaF, AbaQ, AmvA | tet(B), tetR | abeS | abeM | mphE | msrE | catI | | LpsB | MDR |
| AbHTMGP-5566 | AbaF, AbaQ, AmvA | | abeS | abeM | mphE | msrE | | | LpsB | MDR |
| AbHTMGP-24 | AbaF, AbaQ, AmvA | tet(B), tetR | abeS | abeM | mphE | msrE | catI | | LpsB | MDR |
| AbHTMGP-250 | AbaF, AbaQ, AmvA | tet(B), tetR | abeS | abeM | mphE | msrE | catI | | LpsB | MDR |
| AbHTMGP-313 | AbaF, AbaQ, AmvA | tet(B), tetR | abeS | abeM | mphE | msrE | catI | | LpsB | MDR |
| AbHTMGP-391 | AbaF, AbaQ, AmvA | tet(B), tetR | abeS | abeM | mphE | msrE | catI | | LpsB | MDR |
| AbHTMGP-388 | AbaF, AbaQ, AmvA | | abeS | abeM | mphE | msrE | | | LpsB | MDR |
| AbHTMGP-545 | AbaF, AbaQ, AmvA | tet(B), tetR | abeS | abeM | mphE | msrE | catI | | LpsB | MDR |
| AbHTMGP-626 | AbaF, AbaQ, AmvA | tet(B), tetR | abeS | abeM | mphE | msrE | catI | | LpsB | MDR |
| AbHTMGP-863 | AbaQ, AmvA | tet(B), tetR | abeS | abeM | | | | | LpsB | MDR |
| AbHTMGP-917 | AbaF, AbaQ, AmvA | tet(B), tetR | abeS | abeM | | | | | LpsB | MDR |
| AbHTMGP-948 | AbaF, AbaQ, AmvA | tet(B), tetR | abeS | abeM | mphE | msrE | catI | | LpsB | MDR |
| AbHTMGP-949 | AbaF, AbaQ, AmvA | tet(B), tetR | abeS | abeM | | | | | LpsB | MDR |
| AbHTMGP-1125 | AbaF, AbaQ, AmvA | | abeS | abeM | mphE | msrE | | | LpsB | MDR |

| NCBI | SAM | TZP | CAZ | FEP | CTX | CRO | IPM | MEM | GM | AN | TE | CIP | LVX | Tigecycline µg/mL | Colistin µg/mL |
|---|---|---|---|---|---|---|---|---|---|---|---|---|---|---|---|
| AbHTMGP-1608 | R | R | R | R | R | R | R | R | R | R | S | R | R | 1 | 0.5 |
| AbHTMGP-1980 | R | R | I | R | R | R | R | R | R | R | S | R | R | 1 | 0.5 |
| AbHTMGP-2026 | R | R | S | R | R | R | R | R | I | R | S | R | R | 2 | 0.5 |
| AbHTMGP-2007 | R | R | I | R | R | R | R | R | R | R | S | R | R | 2 | 0.5 |
| AbHTMGP-2027 | R | R | I | R | R | R | R | R | R | R | S | R | R | 2 | 0.5 |
| AbHTMGP-2600 | R | R | I | R | R | R | R | R | R | R | S | R | R | 2 | 0.5 |
| AbHTMGP-2689 | S | S | S | S | I | I | S | S | S | S | S | S | S | 0.5 | 0.25 |
| AbHTMGP-2693 | R | R | I | R | R | R | R | R | R | R | S | R | R | 2 | 0.5 |
| AbHTMGP-2957 | R | R | R | R | R | R | R | R | R | R | S | R | R | 2 | 0.5 |
| AbHTMGP-3088 | R | R | R | R | R | R | R | R | R | S | S | R | R | 2 | 1.0 |
| AbHTMGP-3133 | R | R | R | R | R | R | R | R | R | S | S | R | R | 2 | 0.5 |
| AbHTMGP-3145 | R | R | R | R | R | R | R | R | R | R | S | R | R | 1 | 0.5 |
| AbHTMGP-3895 | R | R | R | R | R | R | R | R | R | R | S | R | R | 0.5 | 0.5 |
| AbHTMGP-3910 | R | R | R | R | R | R | R | R | I | S | S | R | R | 2 | 1.0 |
| AbHTMGP-3916 | R | R | R | R | R | R | R | R | R | R | S | R | R | 0.5 | 0.5 |
| AbHTMGP-3971 | R | R | R | R | R | R | R | R | R | R | R | R | R | 1 | 1.0 |
| AbHTMGP-4173 | R | R | R | R | R | R | R | R | R | R | I | R | R | 2 | 1.0 |
| AbHTMGP-4330 | R | R | R | R | R | R | R | R | R | R | I | R | R | 2 | 0.5 |
| AbHTMGP-4473 | R | R | I | R | R | R | R | R | R | R | I | R | R | 4 | 0.5 |
| AbHTMGP-4696 | R | R | R | R | R | R | R | R | R | R | R | R | R | 2 | 1.0 |
| AbHTMGP-4701 | R | R | R | R | R | R | R | R | R | R | R | R | R | 2 | 0.5 |
| AbHTMGP-4826 | R | R | I | R | R | R | R | R | R | R | R | R | R | 2 | 0.5 |
| AbHTMGP-4741 | R | R | I | R | R | R | R | R | R | R | S | R | R | 8 | 0.5 |
| AbHTMGP-4890 | R | R | R | R | R | R | R | R | R | R | I | R | R | 2 | 0.5 |
| AbHTMGP-4888 | R | R | R | R | R | R | R | R | R | R | R | R | R | 2 | 0.5 |
| AbHTMGP-4880 | R | R | S | R | R | R | R | R | R | S | R | R | R | 1 | 1.0 |
| AbHTMGP-4898 | R | R | R | R | R | R | R | R | R | R | R | R | R | 2 | 0.5 |
| AbHTMGP-4911 | R | R | R | R | R | R | R | R | R | R | R | R | R | 4 | 0.5 |
| AbHTMGP-5204 | R | R | R | R | R | R | R | R | R | R | I | R | R | 2 | 1.0 |
| AbHTMGP-5573 | R | R | R | R | R | R | R | R | R | R | R | R | R | 2 | 0.5 |
| AbHTMGP-5566 | R | R | R | R | R | R | R | R | R | R | R | R | R | 4 | 1.0 |
| AbHTMGP-24 | R | R | R | R | R | R | R | R | R | R | R | R | R | 4 | 1.0 |
| AbHTMGP-250 | R | R | R | R | R | R | R | R | R | R | R | R | R | 1 | 0.5 |
| AbHTMGP-313 | R | R | R | R | R | R | R | R | R | R | R | R | R | 2 | 0.5 |
| AbHTMGP-391 | R | R | R | R | R | R | R | R | R | R | R | R | R | 4 | 1.0 |
| AbHTMGP-388 | R | R | R | R | R | R | R | R | R | S | S | R | R | 0.5 | 0.5 |
| AbHTMGP-545 | R | R | R | R | R | R | R | R | R | R | R | R | R | 4 | 0.5 |
| AbHTMGP-626 | R | R | R | R | R | R | R | R | R | R | R | R | R | 4 | 0.5 |
| AbHTMGP-863 | I | R | R | R | R | R | R | R | R | R | S | R | R | 2 | 2.0 |
| AbHTMGP-917 | R | R | R | R | R | R | R | R | R | R | S | R | R | 2 | 0.5 |
| AbHTMGP-948 | R | R | R | R | R | R | R | R | R | R | R | R | R | 4 | 0.5 |
| AbHTMGP-949 | R | R | R | R | R | R | R | R | R | R | I | R | R | 2 | 2.0 |
| AbHTMGP-1125 | R | R | R | R | R | R | R | R | R | R | I | R | R | 4 | 0.5 |

Supplementary Table 2

**Supplemenntary Table 2. *In silico* prediction and typing of potential plasmids.**

| Strains | Contigs with plasmid prediction | Contigs meeting criteria | Plasmid typing by Hamindian's database | Plasmid typing by Bertini's database, Rep group (Rep superfamily)& |
|---|---|---|---|---|
| AbHTMGP-863 | 12 | 1 | p(REP3) | (Rep-3) |
| AbHTMGP-4880 | 2 | 0 | - | |
| AbHTMGP-4911 | 3 | 0 | - | |
| AbHTMGP-388 | 3 | 1 | p(MOBQ ,REP3) | GR4 (Rep-3) |
| AbHTMGP-1125 | 7 | 1 | p(MOBQ ,REP3) | GR4 (Rep-3) |
| AbHTMGP-2957 | 3 | 1 | p(MOBQ ,REP3) | GR4 (Rep-3) |
| AbHTMGP-3145 | 13 | 1 | p(MOBQ ,REP3) | GR4 (Rep-3) |
| AbHTMGP-5566 | 2 | 1 | p(MOBQ ,REP3) | GR4 (Rep-3) |
| AbHTMGP-917 | 15 | 1 | p(MOBP), p(Un) | |
| AbHTMGP-1608 | 2 | 1 | p(MOBP) | |
| AbHTMGP-2026 | 5 | 1 | p(MOBP) | |
| AbHTMGP-2027 | 2 | 1 | p(MOBP) | |
| AbHTMGP-3910 | 4 | 1 | p(MOBP) | |
| AbHTMGP-3971 | 10 | 1 | p(MOBP) | |
| AbHTMGP-4330 | 13 | 1 | p(MOBP) | |
| AbHTMGP-1980 | 6 | 2 | p(MOBQ ,REP3), p(MOBQ) | GR4 (Rep-3) |
| AbHTMGP-2007 | 6 | 2 | p(MOBQ ,REP3), p(MOBQ) | GR4 (Rep-3) |
| AbHTMGP-2600 | 7 | 2 | p(MOBQ ,REP3), p(MOBQ) | GR4 (Rep-3) |
| AbHTMGP-2693 | 6 | 2 | p(MOBQ ,REP3), p(MOBQ) | GR4 (Rep-3) |
| AbHTMGP-4473 | 6 | 2 | p(MOBQ ,REP3), p(MOBQ) | GR4 (Rep-3) |
| AbHTMGP-4741 | 7 | 2 | p(MOBQ ,REP3), p(MOBQ) | GR4 (Rep-3) |
| AbHTMGP-4826 | 7 | 2 | p(MOBQ ,REP3), p(MOBQ) | GR4 (Rep-3) |
| AbHTMGP-313 | 6 | 1 | p(Un)[c] | (Rep-3) |
| AbHTMGP-391 | 6 | 1 | p(Un)[c] | (Rep-3) |
| AbHTMGP-545 | 12 | 1 | p(Un)[c] | (Rep-3) |
| AbHTMGP-626 | 12 | 1 | p(Un)[c] | (Rep-3) |
| AbHTMGP-948 | 12 | 1 | p(Un)[c] | |
| AbHTMGP-4696 | 14 | 1 | p(Un)[c] | (Rep-3) |
| AbHTMGP-4701 | 6 | 1 | p(Un)[c] | (Rep-3) |
| AbHTMGP-4898 | 7 | 1 | p(Un)[c] | (Rep-3) |
| AbHTMGP-24 | 12 | 1 | p(REP3)[c] | |
| AbHTMGP-250 | 12 | 1 | p(REP3)[c] | |
| AbHTMGP-2689 | 1 | 1 | p(REP3)[c] | (Rep-3) |
| AbHTMGP-4888 | 12 | 1 | p(REP3)[c] | |
| AbHTMGP-5573 | 12 | 1 | p(REP3)[c] | |
| AbHTMGP-949 | 17 | 3 | p(MOBP), p(REP3), p(Un)[c] | (Rep-3) |
| AbHTMGP-3133 | 18 | 2 | p(MOBP, REP3)[c], p(REP3) | |
| AbHTMGP-3088 | 16 | 2 | p(MOBQ, REP3)[c], p(MOBP) | GR12 (Rep-3) |
| AbHTMGP-3895 | 6 | 2 | p(MOBQ, REP3)[c], p(MOBP) | GR4 (Rep-3) |
| AbHTMGP-3916 | 13 | 2 | p(MOBQ, REP3)[c], p(MOBP) | GR4 (Rep-3) |
| AbHTMGP-5204 | 15 | 2 | p(MOBQ, REP3)[c], p(MOBP) | GR4 (Rep-3) |
| AbHTMGP-4890 | 3 | 3 | p(MOBQ, REP3)[c], p(MOBP)[c], p(Un)[c] | GR4 (Rep-3) |
| AbHTMGP-4173 | 7 | 4 | MOBQ, REP3)[c], p(MOBQ), p(REP3), p(U | (Rep-3) |

Relaxase typing (MOB) or Replicase typing (R) or Circularity status ([c]); Un, unclassified

& in some instances given that the replicase names in Bertini's paper were defunc, we employed the pfam domain described in the paper but just focusing on proteins decribed in *A. baumannii*
